# Isotropic myosin-generated tissue tension is required for the dynamic orientation of the mitotic spindle

**DOI:** 10.1101/328161

**Authors:** Maxine SY Lam, Ana Lisica, Nitya Ramkumar, Yanlan Mao, Guillaume Charras, Buzz Baum

## Abstract

The ability of epithelial cells to divide along their long cell axis, known as “Hertwig’s rule”, has been proposed to play an important and wide-ranging role in homogenising epithelial cell packing during tissue development and homeostasis. Since the position of the anaphase spindle defines the division plane, how divisions are oriented requires an understanding of the mechanisms that position the mitotic spindle. While many of the molecules required to orient the mitotic spindle have been identified in genetic screens, the mechanisms by which spindles read and align with the long cell axis remain poorly understood. Here, in exploring the dynamics of spindle orientation in mechanically distinct regions of the fly notum, we find that the ability of cells to properly orient their divisions depends both on cortical cues and on local tissue tension. Thus, spindles align with the long cell axis in tissues in which isotropic tension is elevated, but fail to do so in elongated cells within the crowded midline, where tension is low. Importantly, these region-specific differences in spindle behaviour can be reversed by decreasing or increasing the activity of non-muscle Myosin II. In addition, spindles in a tissue experiencing isotropic stress fail to align with the long cell axis if cells are mechanically isolated from their neighbours. These data lead us to propose that isotropic tension is required within an epithelium to provide cells with a mechanically stable substrate upon which localised cortical Dynein can pull on astral microtubules to orient the spindle.

## INTRODUCTION

A typical interphase epithelial cell has a well-defined apico-basal polarity that is maintained by mutually antagonistic interactions between conserved sets of polarity proteins positioned at different sites along the cell’s apico-basal axis [1–3]. To maintain and transmit this polarity axis during cell division so that each daughter cell inherits an apical and basal domain, the mitotic spindle must be properly aligned with the plane of the tissue [4,5] as it defines the cell division plane at anaphase [6,7]. In many systems, positioning the spindle in the plane of the tissue, or *z*-positioning, is important since mis-orientation can result in defects in epithelial organisation [8,9] and the loss of cells from the epithelium [10–13]. Within the plane of the tissue, *xy*-positioning of the spindle is also important. Spindles positioned at the centre of the cell allow for the equal distribution of cellular material between daughter cells during a symmetric division [14,15], while spindle alignment with the long cell axis helps to limit cell elongation. The latter is often known as the long-axis or Hertwig’s rule [5,16–20], and has been suggested to play important roles in maintaining homeostatic cell packing and in aiding the relaxation of tissues subjected to mechanical strain [17,20–23]. However, the mechanisms that enable spindles to detect and orient with the long cell axis in the plane of the epithelium remain poorly understood.

Spindles are positioned by forces acting on astral microtubules (MTs) emanating from the centrosomes at opposing spindle poles. In most cases Dynein-based pulling forces appear to play a dominant role in positioning the mitotic spindle [23–28]. In mitotic epithelial cells, this Dynein is associated with Mud/NuMA/LIN-5 [5,29] at the cell cortex [5,25,30,31]. The active Dynein complex anchored to the cortex then walks towards the minus-ends of astral microtubules, effectively ‘reeling’ in the centrosome to reposition the spindle. Recently in the fly notum, Mud/NuMA was also found enriched at tricellular junctions (TCJs, where 3 or more cells come into contact) in interphase and mitotic cells [23]. This localisation was found to depend on the TCJ proteins, Dlg (Discs large) and Gli (Gliotactin), which the authors showed are also required for mitotic spindle orientation in this tissue [23]. These observations led the authors to propose that TCJ-localised Mud/NuMA links interphase cell shape to mitotic spindle orientation [23]. A theoretical model based on TCJs was able to predict experimental spindle orientation to within 30° [23]. While impressive, this suggests that the system is noisy. Moreover, Mud/NuMA is not found enriched at TCJs in other tissues and model systems [22,29,32,33], raising the possibility that TCJ-localised Mud/NuMA may only be one component of the system in the fly notum, and that additional mechanisms might be involved in spindle orientation more generally [34].

One such factor may be mechanical tension. Extrinsic mechanical cues can orient mitotic spindles in isolated cells in culture, where spindles position according to the pattern of actin-based retraction fibres that connect cells to the substrate [35–37], and spindles are able to rotate to align with a tension axis independent of cell shape in prometaphase [38]. In epithelial tissues, spindles have also been shown to orient along a stretch axis [20–22,39,40]. However, it remains unclear whether spindles in epithelia orient in response to stretch by directly sensing and aligning with the imposed tension axis and/or by aligning with the long cell axis and/or to the position of TCJs, since all of these cues tend to align in the same direction [20–22,39,40].

Here, to determine the relative importance of shape and tension in spindle orientation, we compared the dynamics of spindle orientation within the distinct mechanical environments present in the developing fly notum midline. The notum is a single-layered epithelium that lies at the surface of developing fly pupa, making it an excellent system in which to combine genetics and live cell imaging in the study of cell division [41–45]. Previous work in the notum has demonstrated that cells within the tissue midline (ML) experience significant crowding but little to no tension, despite being very elongated [46]. By contrast, cells outside the midline (OML) are less elongated but experience elevated levels of isotropic tissue tension [46,47]. This decoupling of shape and tension makes the notum an ideal system in which to determine their relative contributions to spindle orientation. Strikingly, our analysis revealed that mis-oriented spindles in the crowded midline of the wildtype tissue could not re-orient to obey the long-axis rule – necessitating the use of alternative mechanisms of cells packing refinement [46]. Importantly, the effects of endogenous high/low tissue tension could be recapitulated by modifying tissue tension through the modulation of non-muscle Myosin II activity or laser ablation. Furthermore, we show that these differences in spindle re-orientation result from differences in the persistence of spindle rotation towards the long cell axis, which depend in turn on the local levels of isotropic tissue tension. Based on these data, we propose a model for epithelial spindle orientation in which tissue tension promotes the effective rotation of spindles towards the long cell axis by providing a mechanically stable substrate upon which localised cortical Dynein can pull.

## RESULTS

### Cells in the fly notum midline fail to divide along to their long cell axis

Cells in the central part of the notum undergo a single round of division between 15-19h after pupariation (AP) [42]. At the start of this period, cells in the tissue midline appear very elongated (Fig S1A-B). Nevertheless, because of tissue crowding, these cells experience little isotropic tension, as measured by laser ablations of adherens junctions [46]. By contrast, cells outside of the midline are subjected to elevated levels of isotropic tension [46,47], despite being less elongated (Fig S1A-B). By 19h AP, however, when the round of division is complete, cell shapes, cell packing and tissue mechanics have become much more homogenous across the entire tissue (Fig S1A) - a process we have termed tissue refinement [46,47]. We exploited these regional mechanical differences to investigate how tissue tension affects cell division to the long axis to aid this process.

To follow spindle orientation relative to cell shape over time, Tubulin-mCherry was used to visualize the spindle or Centrosomin-RFP to mark spindle poles, while Spider-GFP labelled the nuclear envelope and the cell membrane (Fig 1A). By tracking both spindle poles in 3D, we found that the vast majority of mitotic spindles were positioned within the plane of the epithelium at NEB (measured by loss of the nuclear envelope marker) or shortly thereafter, as indicated by the localisation of both centrosomes within a 1.5μm height of one another and often within a single imaging plane (Fig 1A, Fig S2). Spindles then remained within the tissue plane throughout mitosis, even though they continued to rotate. These rotational movements in *x-y* were significant (Fig S2), with spindles in cells both inside and outside of the midline undergoing net rotations (θ_displacement._ = angle between anaphase and NEB orientation) of ~45° (Fig S1C) and cumulative rotations (θ_cumulative rotation_ = sum of change in rotation per minute) of >90° (Fig S1D). Since the average θ_cumulative rotation_ was approximately twice that of the average θ_displacement_, these results indicate that spindle orientation is a dynamic and noisy process irrespective of local differences in tissue tension.

**Fig 1:**
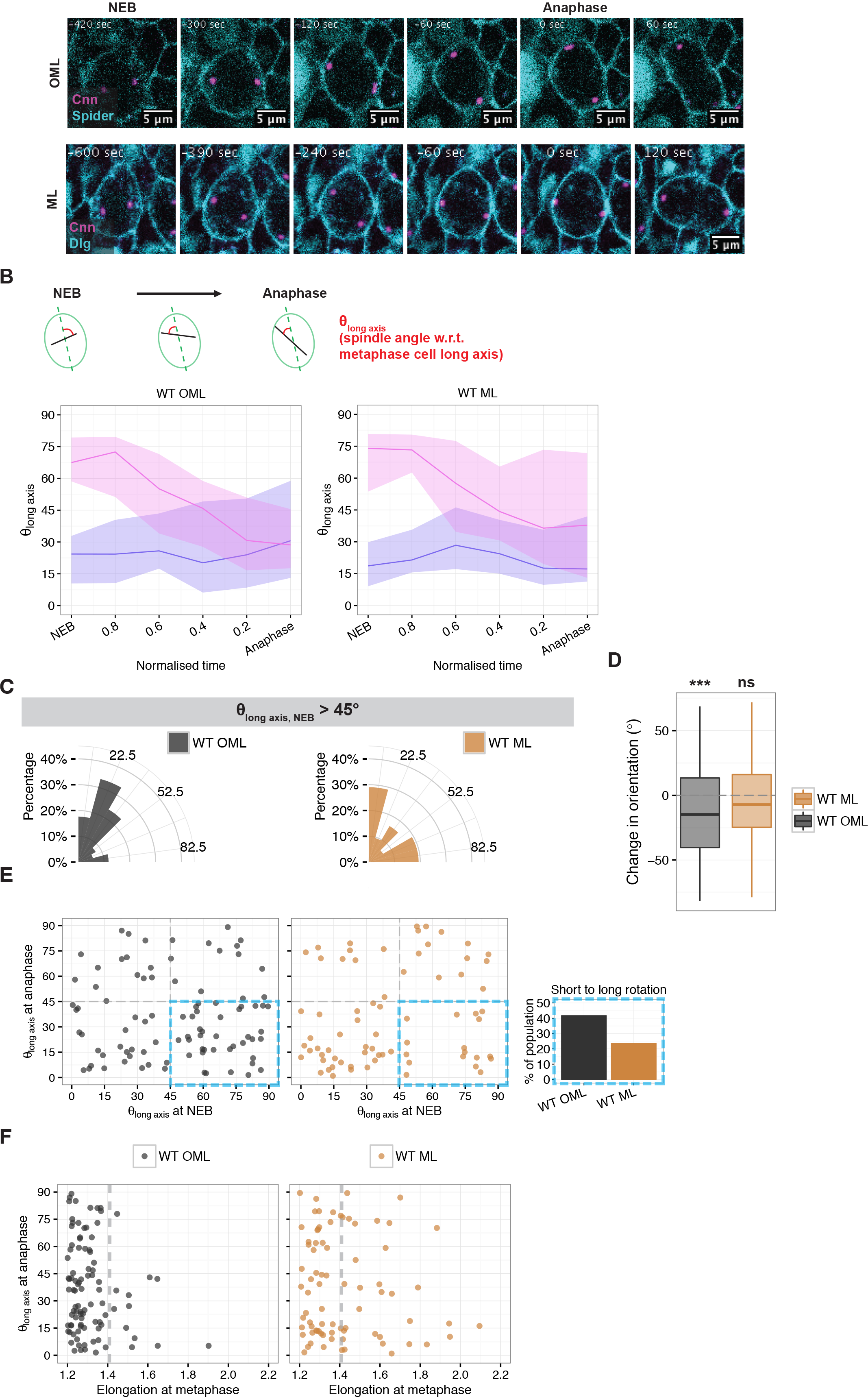
Cells in the Drosophila notum midline fail to divide along their long cell axis. (Related to Fig S1 and S2) **A**: Example OML and ML cells during mitosis and diagram of spindle rotation analysis. Cell membranes including labelled with Spider-GFP (cyan) and spindle centrosomes are labelled with Centrosomin-RFP (magenta). θ_long axis_ was defined as the angle between the spindle and the long axis of the cell at metaphase. **B**: θ_long axis_ from NEB to anaphase for OML and ML spindles. Data is shown for subpopulation of spindles that are ≥45° (pink line) or <45° (purple line) at NEB. Lines indicate median values and shaded regions indicate interquartile range. **C**: θ_long axis_ at anaphase for OML and ML where θ_long axis_ at NEB >45°. The distribution of θ_long axis_ for OML spindles is smaller and closer to 0° than that for ML spindles. **D**: Change in orientation for OML and ML spindles. Change in orientation was calculated as (θ_long axis_ at anaphase- θ_long axis_ at NEB), and data was tested in a one-tail non-parametric test (less than) against 0 as a null hypothesis. Change in orientation was significantly <0 for OML spindles, indicating rotation towards the long axis (−12.0±3.8, p=0.0014, n=91). Change in orientation was no significantly <0 for ML spindles (−5.3±4.1, p=0.099, n=72). **E**: Orientation of spindles at anaphase against at NEB for OML and ML spindles. OML cells have more spindles that are oriented >45° at NEB but <45° at anaphase, compared to ML cells (blue box). **F**: Spindle orientation at anaphase against cell elongation at metaphase for OML and ML cells. Almost all OML cells at higher elongations (>1.4) had an orientation of <45° at anaphase while ML cells with higher elongations often had an orientation of >45° at anaphase.

To investigate whether these rotational movements were accompanied by the orientation of spindles relative to the long axis, we tracked the orientation of spindles from NEB to anaphase, beginning with cells outside of the midline. Spindle orientation (θ_long axis_) was defined as the angle between the spindle and the major axis of an ellipse fitted to the cell shape at metaphase (Fig 1B). Shortly after NEB, θ_long axis_ appeared random. However, by anaphase, θ_long axis_ was within 45° of the long axis for majority of spindles (Fig 1C). Further analysis of θ_long axis_ over time revealed that the change in spindle orientation depended on the orientation at NEB relative to the long cell axis. Thus, spindles that were mis-oriented at NEB (θ_long axis_ >45° at NEB, Fig 1B [pink line], 1E) underwent persistent, corrective rotational movements towards the long axis during the course of mitosis, while spindles that were already well-oriented at NEB (θ_long axis_ <45° at NEB, Fig 1B [purple line]) did not undergo a significant change in their orientation with time. In combination, these two effects led to an overall change in spindle orientation towards the long cell axis during the course of mitosis (Fig 1D). Thus, in cells outside of the notum midline, spindles obey the long axis rule, as previously published [23], by actively rotating towards the cell long axis.

However, this was not the case within the midline. Here, a significant proportion of spindles with θ_long axis_ >45° at NEB failed to rotate towards the long axis (Fig 1B, 1C, 1E), even though they rotated a similar amount compared to cells outside the midline (Fig S1C-F). As a result, there was no overall change in spindle orientation towards the long axis (Fig 1D). This was surprising because current models of spindle orientation predict that long-axis orientation is more accurate as cell shape becomes more anisotropic, and these cells are extremely elongated during mitosis (Fig S1A). However, the correlation between aspect ratio and spindle orientation was extremely poor for cells in the midline (Coefficient = −1.44 × 10^−3^, p = 0.14) (Fig 1E). In addition, ~31% (11/35) of the spindles in ML cells with an AR >1.4 failed to align to within 45° of the long cell axis (Fig 1F). By contrast, for cells outside of the midline there was a relatively good correlation between aspect ratio and spindle alignment (Coefficient = −8.60 × 10^−4^, p = 0.07) (Fig 1E), and, within this region, only ~8% (1/12) spindles in cells with an AR >1.4 failed to align to within 45° of the long cell axis (Fig 1F). These data reveal surprising, region-specific differences in the ability of spindles to align with the long cell axis.

Because Mud/NuMA, a conserved protein that has been shown to play an important role in spindle orientation in many systems (Morin & Bellaïche 2011), has been shown to localise to TCJs [23], we also considered the possibility that spindle mis-orientation could be explained by an unusual distribution of TCJs around cells in the midline. However, TCJ polarity remained closely aligned with the cell long axis in the midline, even in cells that failed to orient their spindles along the long cell axis (Fig S1H, S2B, S2D). Thus, spindles in the midline of the notum are unable to read their cell long axis. This makes the notum midline one of the few examples of a tissue in which symmetric divisions disobey the long-axis rule (reviewed in [5]).

### Cortical forces acting on astral microtubules are required for dynamic spindle positioning

Since spindles in cells outside of the midline are able to rotate to align with the long cell axis, we were able to use this region to explore the mechansims involved in spindle orientation in more detail. Previous work in the notum has suggested roles for Dynein, Mud/NuMA and Dlg in the orientation of spindles to the long cell axis [23]. We therefore tested the role of Mud/NuMA outside of the midline using Mud RNAi (Mud-IR). As previously reported [23], Mud-IR resulted in varying degrees of *z*-positioning defects. Despite this, ~50% of spindles in Mud-IR cells aligned well with the plane of the epithelium (measured as a difference in pole to pole height of ≤1.5μm) (Fig 2A). Within these cells, we observed very little spindle rotation over time (Fig 2A-D). Accordingly, there was no global change in spindle orientation in Mud-IR cells from NEB to anaphase (Fig 2E). Since cortical forces pulling on astral microtubules can also centre spindles [18,24,48], we also examined the impact of Mud/NuMA on spindle centring in this system. Strikingly, mitotic spindles were closer to the cell centre in metaphase Mud-IR cells than they were in WT cells (WT OML: 1.00±0.07μm, n=58; Mud-IR OML: 0.59±0.03μm, n=89. Fig 2F). Thus, the Mud/NuMA depedent forces that orient the spindle also have a tendency to pull the spindle off-centre.

**Fig 2:**
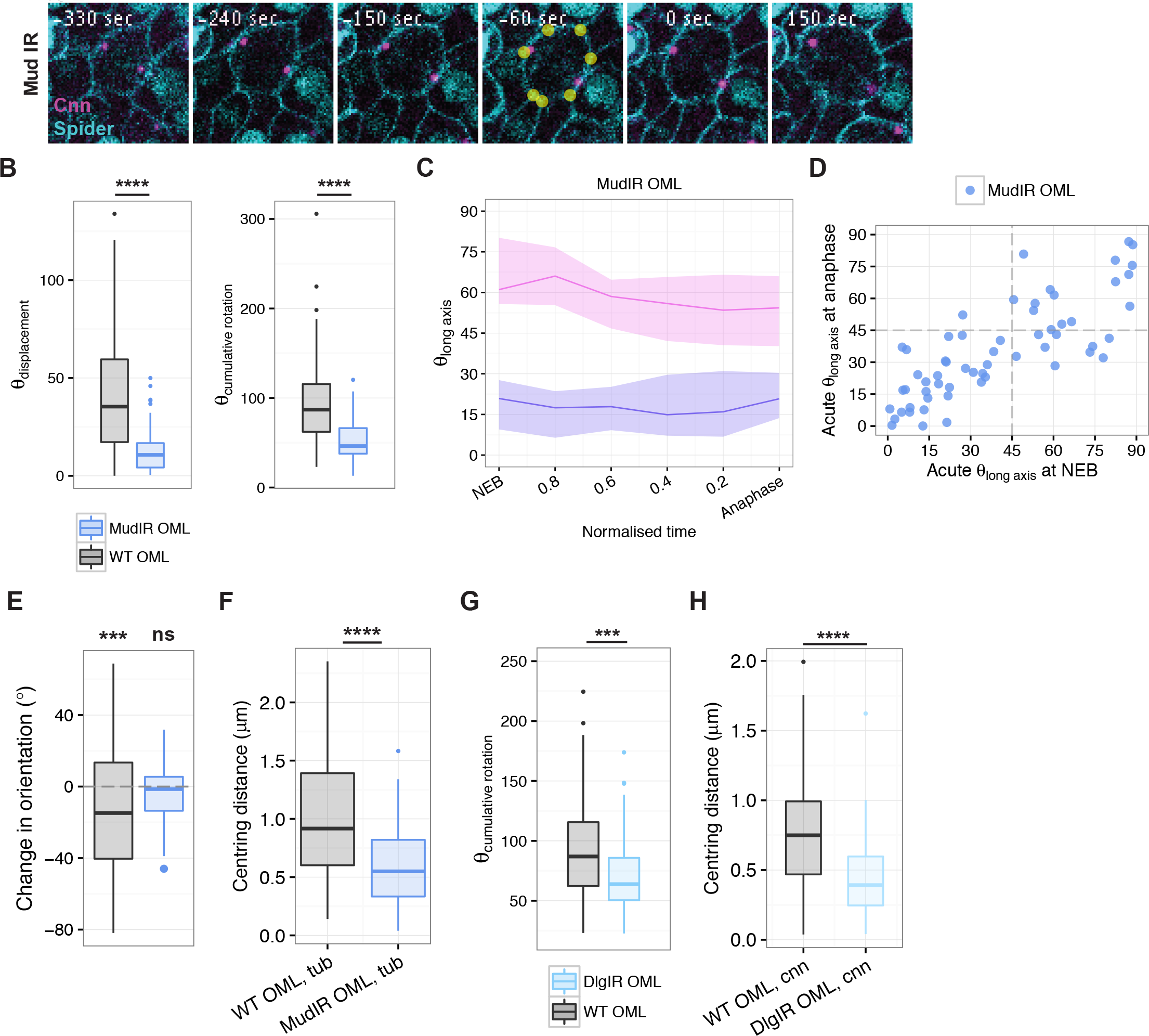
Cortical forces acting on astral microtubules are required for dynamic spindle positioning. (Related to Fig S3) **A**: Example Mud-IR cell with no severe *z*-positioning defect but severe *xy*-positiong defect during mitosis. Yellow dots indicate TCJs. **B**: θ_displacement_ and θ_cumulative rotation_ for WT and Mud-IR OML spindles. θ_displacement_ and θ_cumulative rotation_ was dramatically reduced in Mud-IR cells (θ_displacement_ – WT: 42.76°±3.48°, n=91; Mud-IR: 13.15°±1.58°, n= 59; p=1.432×10^−9^. θ_cumulative rotation_ – WT: 94.55°±4.87°, n=91; Mud-IR: 53.77°±2.93°, n=59; p<2.2×10^−16^). **C**: θ_long axis_ from NEB to anaphase for Mud-IR OML spindles. Data is shown for subpopulation of spindles that are ≥45° (pink line) or <45° (purple line) at NEB. Lines indicate median values and shaded regions indicate interquartile range. **D**: Orientation of spindles at anaphase against at NEB for Mud-IR OML spindles. Majority of spindles have little change in orientation from NEB to anaphase. **E**: Change in orientation for WT and Mud-IR OML spindles. Change in orientation was ≈0 for Mud-IR OML spindles (−3.92±2.18, p=0.088, n=59). **F**: Centring distance for WT and MudIR OML spindles labelled with Tubulin-mCherry. Centring distance was calculated as the linear distance between the centre of the spindle and the centroid of the cell in the same plane at metaphase. MudIR spindles were significantly closer to the centre of the cell compared to WT spindles (WT: 1.00±0.07, n= 58; MudIR: 0.59±0.03, n=89, p=2.77×10^6^). **G**: θ_cumulative rotation_ for WT and DlgIR OML spindles. θ_cumulative rotation_ is significantly reduced in DlgIR OML spindles (DlgIR: 72.68°±5.81°, n=37; p=0.0048). **H**: Centring distance for WT and DlgIR OML spindles labelled with Centrosomin-RFP. DlgIR spindles were significantly closer to the centre of the cell compared to WT spindles (WT: 0.77±0.04, n=84; DlgIR: 0.46±0.04, n=50; p=6.46×10^−06^).

While these data fit with a role for cortical Mud/NuMA in generating the Dynein-mediated pulling forces that act on the spindle, Mud/NuMA is also found localised to spindle poles [49]. Therefore, to specifically compromise cortical pulling forces, we used RNAi to silence expression of the junctional protein Dlg - a protein implicated in the localisation of Mud/NuMA to the cell cortex [23]. To avoid pupal lethality, flies were raised at 18°C to reduce Gal4-mediated Dlg-IR expression. Nevertheless, this was sufficient to significantly reduce the Dlg signal detected by immunostaining (data not shown). This level of Dlg knock-down also resulted in a partial reduction in net rotation (Fig S3B), and a significant reduction in cumulative spindle rotation (Fig 2G), and improved spindle centring (Fig 2H). Finally, since cortical forces act on the spindle via astral microtubules, we looked at spindle positioning in a mutant lacking astral MTs, *Asterless*. Spindle tracking from NEB to anaphase was impossible in this mutant background because of the severe *z*-positioning defects (Fig S3C), but the occasional spindles with a planar orientation at metaphase were more centred than those in WT tissues (*Asl*−/− OML: 0.76 ± 0.05, n=68. Fig S3D).

Taken together, these data support the idea that cortical forces act on astral microtubules to position the spindle during mitosis. More specifically, our analysis indicates that cortical Mud/NuMA is important for spindle rotation that allows for dynamic spindle orientation to the cell long axis; and that cortical forces acting on astral MTs also tend to pull spindles off-centre [25–28,50].

### Myosin activity affects dynamic spindle orientation

Since the active rotation of the spindle towards the long cell axis was much more efficient in cells outside of the midline, where tissue tension is elevated relative to the midline, we tested whether regional differences in tissue tension might explain the regional differences in spindle rotational behaviour. Previous work has suggested that the expression of phospho-dead (SqhAA) and phospho-mimetic (SqhEE) versions of myosin light chain (Spaghetti squash, Sqh) can be used to decrease or increase Myosin function and tissue tension, respectively [47]. Therefore, we used the expression of SqhAA and SqhEE as a first test of the role of tissue tension in spindle orientation.

Strikingly, the expression of SqhAA in cells outside the midline reduced the proportion of spindles able to re-orient towards the long cell axis from ~70% to ~30% (Fig 3A-C, 4A; compare with Fig 1B), but did not induce significant changes in mitotic cell shape or TCJ polarity (Fig 3A). This led to an increase in the proportion of spindles that failed to align with the long cell axis (Fig 3C, 4A; compare with 1C, 1E), and to a reduction in the overall change in spindle orientations from NEB to anaphase relative to the WT control (Fig 3D). SqhAA expression also increased the proportion of metaphase spindles that failed to align with the long axis in very elongated cells (AR >1.4, θ_long axis_ >45° at anaphase) from ~10% to ~30% (12/34, Fig 4B). Thus, SqhAA expression caused spindles in cells outside the midline to behave like spindles in the wildtype midline. Notably, this phenotype was distinct from that observed in Mud-IR cells, since SqhAA expression did not affect net spindle rotation (θ_displacement_) and increased cumulative spindle rotation (θ_cumulative rotation_) (Fig 4C), indicating that spindles can still rotate. Additionally, there was no change in spindle centring in SqhAA-expressing cells outside of the midline compared to WT (Fig S4A). Instead, SqhAA expression reduced rotational persistence of spindles far from the long cell axis (θ_long axis_ >45°), compared to that of WT spindles (Fig 4D). Taken together, these data imply that cortical pulling forces still act on the spindle in tissues expressing SqhAA, but that these are not as effective in directing spindle rotations as they are in cells within tissues experiencing higher levels of isotropic tension.

**Fig 3:**
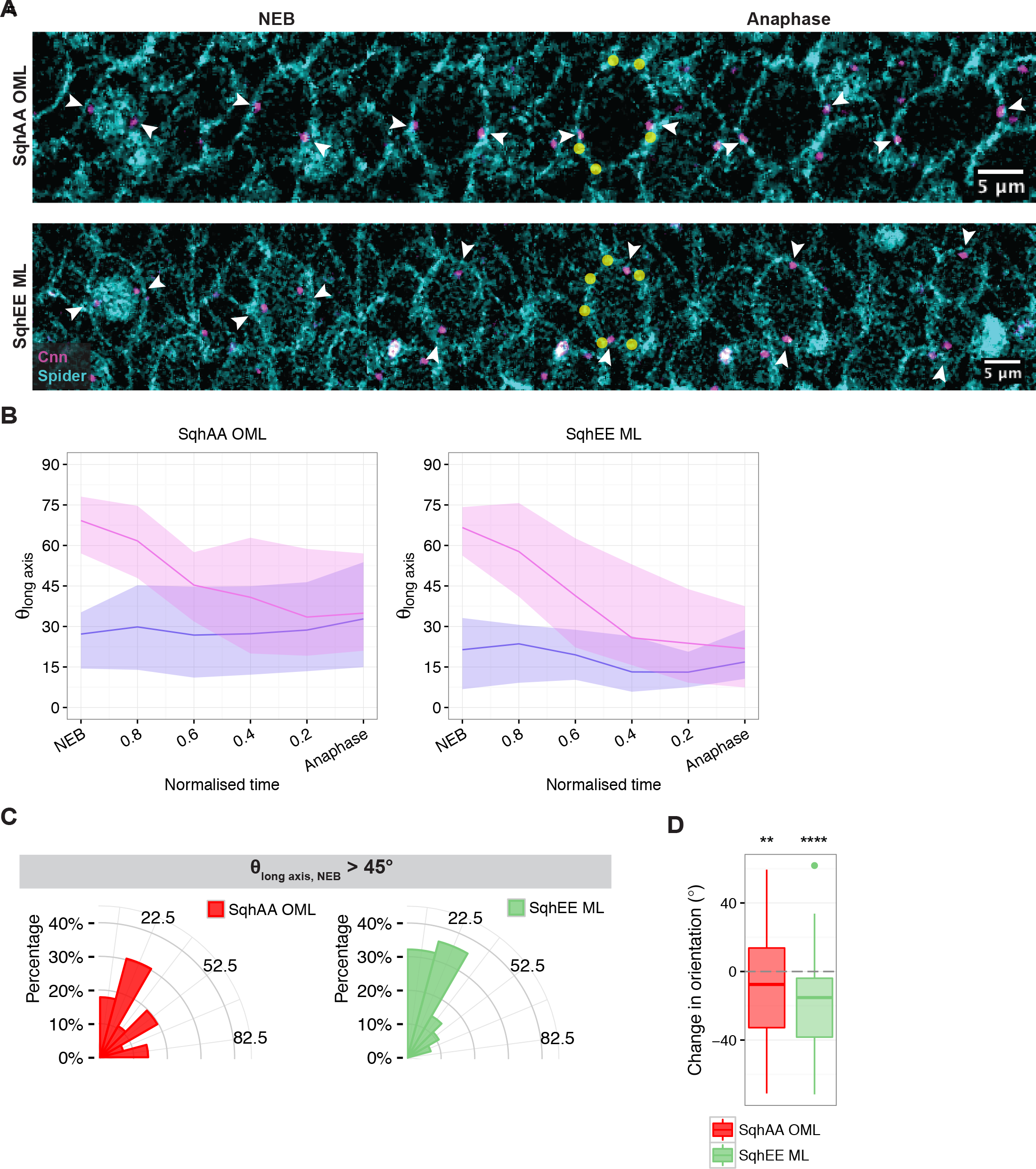
Myosin activity affects dynamic spindle orientation. (Related to Fig S4) **A**: Example SqhAA OML and SqhEE ML cells during mitosis. Cell membranes including nuclear envelope are labelled with Spider-GFP (cyan) and spindle centrosomes are labelled with Centrosomin-RFP (magenta). TCJs are indicated with yellow dots. **B**: θ_long axis_ from NEB to anaphase for SqhAA OML and SqhEE ML spindles. Data is shown for subpopulation of spindles that are ≥45° (pink line) or <45° (purple line) at NEB. Lines indicate median values and shaded regions indicate interquartile range. **C**: θ_long axis_ at anaphase for SqAA OML and SqhEE ML where θ_long axis_ at NEB >45°. The distribution of θ_long axis_ for SqAA OML spindles is wider than for WT OML spindles, the distribution of θ_long axis_ for SqhEE ML spindles is narrower and closer to 0° than WT ML spindles. **D**: Change in orientation for SqhAA OML and SqhEE ML spindles. Change in orientation was <0 for SqhAA OML spindles but this was less significant compared to WT OML spindles (SqhAA OML: −9.09±3.07, p=0.0029, n=109). Meanwhile change in orientation was now significantly <0 for SqhEE ML spindles (SqhEE ML: −19.28±3.78, p=8.15×10^−6^, n=60).

**Fig 4:**
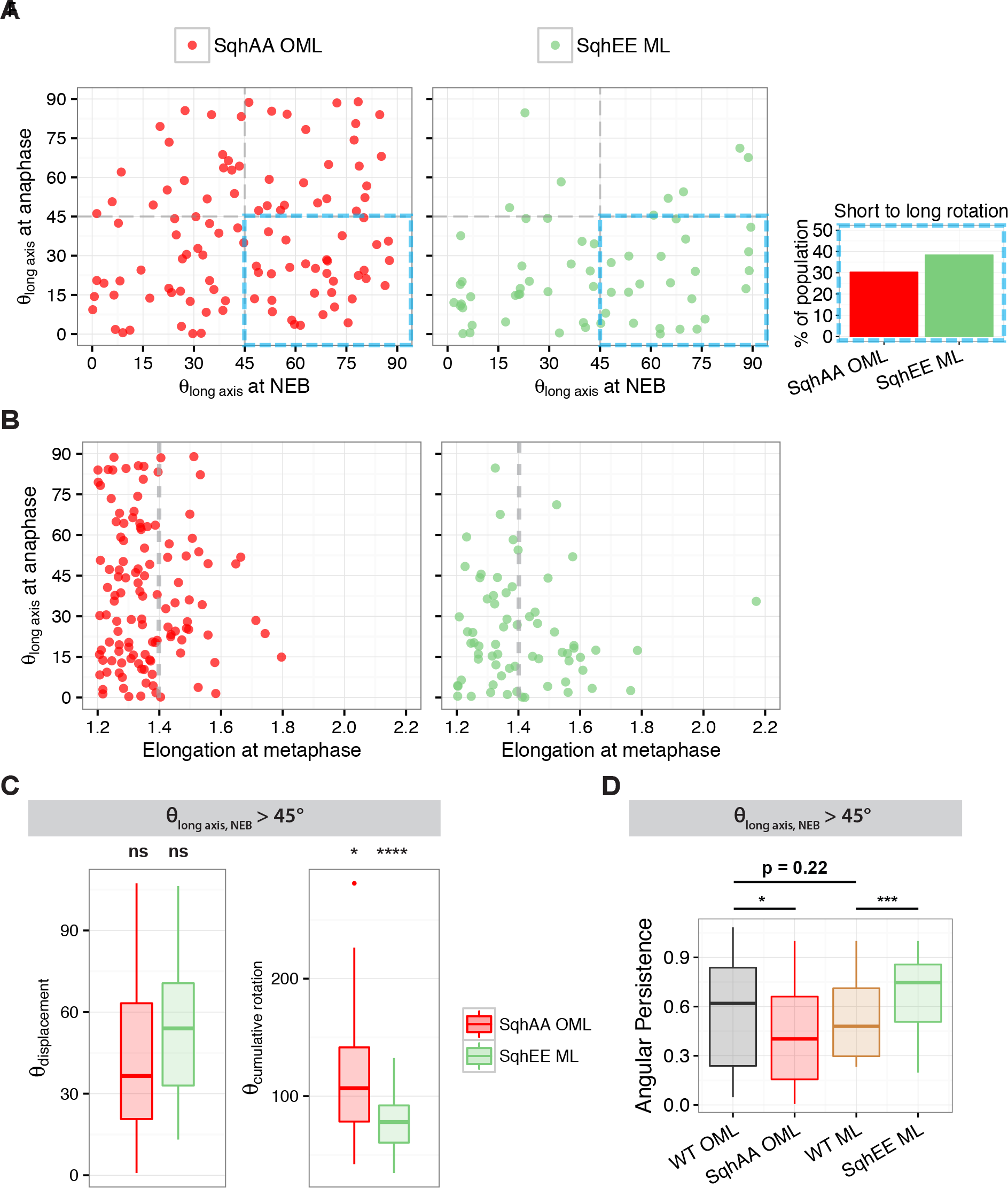
Myosin activity affects directed spindle rotation and orientation. (Related to Fig S4) **A**: Orientation of spindles at anaphase against at NEB for SqhAA OML and SqhEE ML spindles. SqhAA OML cells have less spindles that are oriented >45° at NEB but <45° at anaphase (blue box), compared to WT OML cells. SqhEE ML cells have more spindles that are oriented >45° at NEB but <45° at anaphase (blue box), compared to WT ML cells. **B**: Spindle orientation at anaphase against cell elongation at metaphase for SqhAA OML and SqhEE ML cells. SqhAA OML cells at higher elongations (>1.4) had orientations of >45° at anaphase, while almost all SqhEE ML cells with higher elongations were oriented <45° at anaphase. **C**: θ_displacement_ and θ_cumulative rotation_ for SqhAA OML and SqhEE ML spindles where θ_long axis_ at NEB >45°. θ_displacement_ for SqhAA OML and SqhEE ML was similar to WT OML and ML respectively (SqhAA OML: 42.37°±4.80°, n=36, p=0.24; SqhEE ML: 52.37°±4.55°, n=28, p=0.78). However, θ_cumulative rotation_ was significantly higher for SqhAA OML compared to WT OML (SqhAA OML: 115.26°±8.57°, n=36, p=0.028) and significantly lower for SqhEE ML compared to WT ML spindles (SqhEE ML: 77.54°±3.43°, n=28, p=8.2×10^−4^). **D**: Angular persistence for WT OML, SqhAA OML, WT ML and SqhEE ML spindles that were >45° at NEB. Angular persistence was calculated as the ratio of θ_displacement_/ θ_cumulative rotation_. Angular persistence is similar for WT OML and WT ML spindles (WT OML: 0.56±0.04, n=47; WT ML: 0.50±0.04, n=31; p=0.22), but is reduced in SqhAA OML spindles compared to WT OML spindles (SqhAA OML: 0.44±0.05, n=36, p=0.030) and increased in SqhEE ML spindles compared to WT ML spindles (SqhEE OML: 0.68±0.04, n=28, p=0.0044).

For the converse experiment, we expressed SqhEE to increase tissue tension in the midline. Strikingly, this enhanced the ability of spindles in the midline to rotate towards the long cell axis. Thus, following SqhEE expression, almost all spindles with θ_long axis_ >45° at NEB rotated towards the long axis, such that θ_long axis_ was <45° by anaphase (Fig 3A-C). Additionally, spindles with a θ_long axis_ of <45° at NEB also rotated even more towards the long axis over time (Fig 3B, 4A). This increase in the accuracy of spindle orientation occurred without a corresponding change in the net rotation of spindles (Fig 4C). Instead, the average cumulative rotation (Fig 4C) of spindles was reduced, because of an increased angular persistence of spindle rotation (Fig 4B). Furthermore, in very elongated midline cells (AR >1.4), SqhEE expression also improved spindle orientation, so that only ~8% (2/26) of spindles remained poorly aligned with the long cell axis (θ_long axis_ >45°) at the end of mitosis compared to ~31% in the WT midline cells (Fig 4B).

The ability of SqhEE expression to improve the accuracy of spindle orientation in the midline did not appear to be a peculiarity of the construct or midline cells, since a similar trend was also seen in cells outside of the midline (Fig S4B-E). Here, the already robust rotation of spindles towards the cell long axis was further improved (Fig S4B-E). Moreover, the effects of SqhEE/AA expression were recapitulated by perturbations that affected Rho kinase (ROK), an upstream regulator of myosin activity, although the impact of perturbing ROK activity on spindle rotations was less strong than with SqhAA or SqhEE expression (Fig S5). Thus, silencing Rok using RNAi (ROK-IR) mimicked SqhAA expression, and the rotation of spindles from the short cell axis at NEB to the long cell axis at anaphase was compromised in ROK-IR cells outside of the midline. By contrast, the expression of a constitutively active version of the kinase (ROK-CA), mimicked the effects of SqhEE, and improved spindle rotation to the long cell axis within the midline (Fig S5A-C). Additionally, ROK-CA expression improved spindle orientation in very elongated cells (AR >1.4) within the midline (Fig S5D), whereas spindle orientation in similarly elongated cells outside the midline was compromised by ROK-IR (Fig S5D).

Overall, these data suggest a model in which myosin-mediated tissue tension is involved in dynamic spindle orientation towards the cell long axis, through the promotion of persistent spindle rotation.

### Cell-extrinsic tension is important for dynamic spindle orientation

Finally, to test whether Myosin aids oriented cell division in the notum through its role in tissue tension, as suggested by the experiments above, we used laser ablation as an orthogonal approach to induce a sudden reduction in the isotropic tissue tension surrounding a cell in mitosis. Since such an effect could only be observed in cells in which spindles undergo stereotypic directed rotation towards the cell long axis, tissues expressing SqhEE cells were used for this experiment (Fig 5). Shortly after NEB, SqhEE cells entering mitosis were subjected to a laser cut at the level of adherens junctions to mechanically isolate a cell from the rest of the tissue. To avoid gross changes in mitotic cell shape, these cuts were made at least one cell row away from the mitotic cell (Fig 5A). Under these conditions, junction ablation did not alter cell shape within the medial spindle plane of the tissue, although it led to a small decrease in cell aspect ratio at the apical plane (Fig 5B). Moreover, the cut did not lead to a gross delay in mitosis in the mechanically isolated mitotic cells, and all cells entered anaphase within ~5.5 min of the start of imaging, similar to what is observed without laser ablation (Fig 5C).

**Fig 5:**
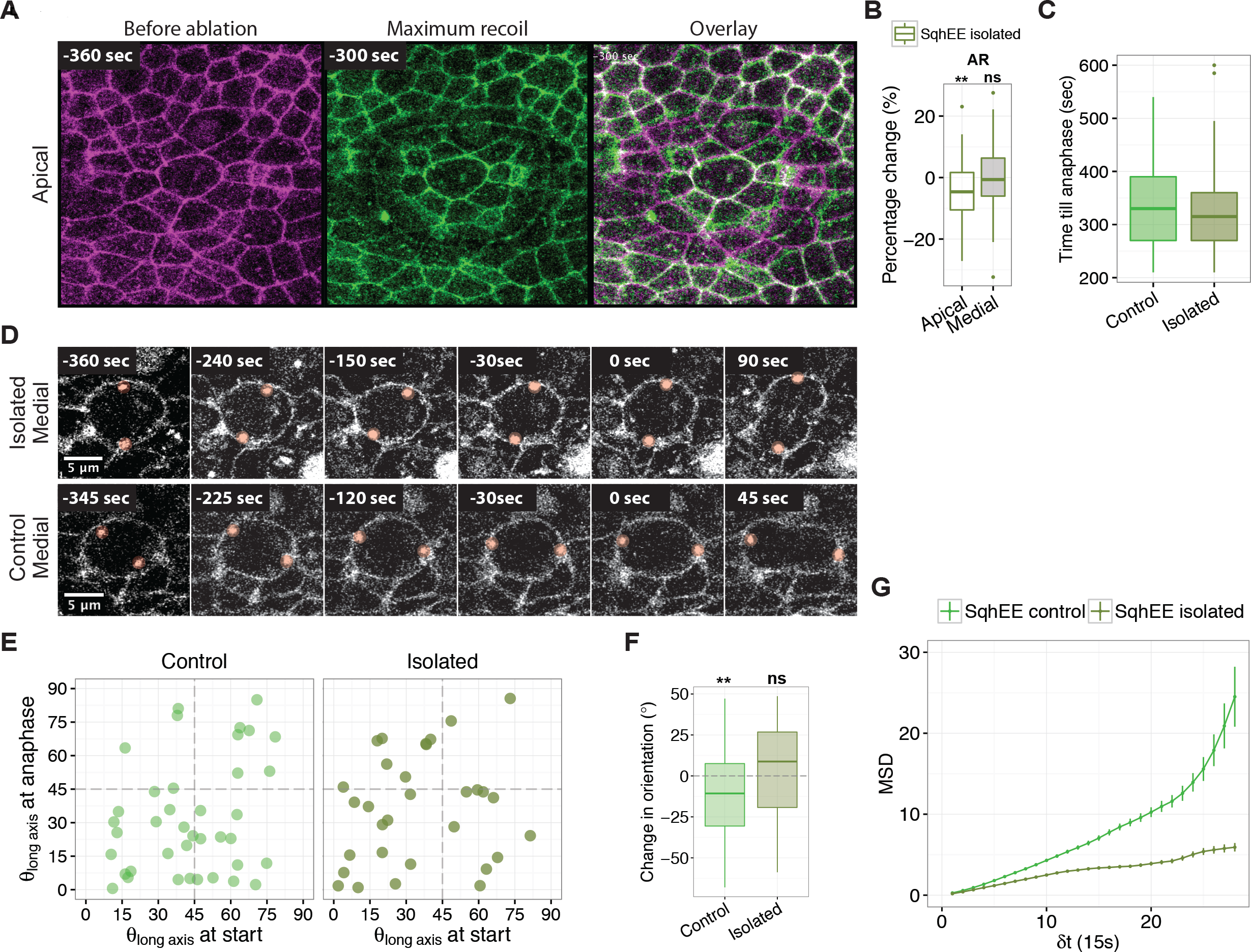
Cell-extrinsic tension is important for dynamic spindle orientation. **A**: Apical surface of a mitotic cell before mechanical isolation by laser ablation (−360 sec till anaphase onset) and after ablation, at the point of maximum recoil of adherens junctions (−300 sec till anaphase onset). Cell membranes were labelled with Spider::GFP. Overlay panel shows extent of recoil in cells neighbouring mitotic cell, but not of the mitotic cell itself. **B**: Change in elongation of mitotic cell at the apical surface and the medial surface (height of spindle) after laser ablation. Cell elongation is reduced at the apical surface but unchanged medially (Apical: −4.28±1.42, p=0.0059, n=53; Medial: −0.53±1.45, p=0.85, n=53). **C**: Time caught in mitosis (from beginning of imaging till anaphase onset) for control and isolated spindles. Time in mitosis was below that of average mitotic time, indicating no significant delays in mitosis. **D**: Montage of mitotic cell viewed at the height of the spindle plane. Cell membranes were labelled with Spider::GFP and spindle centrosomes were labelled with Centrosomin::GFP. **E**: Orientation at anaphase against orientation at start of imaging for control and isolated spindles. **F**: Change in orientation for control and isolated spindles. Change in orientation was calculated as the difference in orientation from the start of imaging till anaphase onset. Change in orientation was significantly <0 for control spindles but not for isolated spindles (Control: −11.45±4.49, p=0.0077, n=39; Isolated: 1.23±5.50, p=0.68, n=31). **G**: Mean Squared Displacement for control and isolated spindles. MSD for control spindles is consistently higher than that for isolated spindles, and increases non-linearly over time.

Within these tissues, changes in spindle orientation were measured at a high frame rate from the beginning of imaging until the onset of anaphase, using spindle movements in SqhEE expressing cells in areas far from the sites of ablation or in tissues where we did not ablate junctions as controls. Since cells about to enter mitosis had to be identified manually before setting up the ablation, we were unable to capture the moment of NEB in these cells. However, we were still able to follow the dynamics of spindle movements. Strikingly, there was a profound difference in spindle movements between the experiment and control. In mechanically isolated SqhEE-expressing cells, spindles failed to rotate towards the long cell axis during the time of imaging, while they did so in the controls (Fig 5D-F). To further investigate the movement of the spindle in these experiments, we calculated the mean-squared displacement (MSD) of each individual centrosome (Fig 4D). The MSD of individual centrosomes per time-step (15 sec) was lower in mechanically isolated cells than it was in the unperturbed tissue (Fig 4G), indicating that directional centrosome movement was in fact reduced after mechanical isolation. Additionally, the MSD over time for mechanically isolated cells was linear, consistent with random motion, while that of unperturbed cells increased non-linearly, indicating active motion. These data lead us to conclude that spindles respond to dynamic changes in the mechanical environment of the mitotic cell and that isotropic tissue tension is required to support effective directed spindle rotation towards the cell long axis.

## DISCUSSION

While an increasing body of research suggests that the forces orienting the mitotic spindles in animal cells are generated by Dynein anchored to the cortex by a set of conserved regulators that include Mud/NuMA and Dlg, the process remains poorly understood. To shed light on the dynamics of the process in the context of an intact epithelium within a developing animal, we imaged spindle orientation in cells of the fly notum expressing probes that label spindle poles, the nuclear envelope and the cell membrane. Since mitotic cell shape has been shown to directly influence spindle positioning [51,52], we focused on cell shape within the same plane as the spindle (a few microns below the apical plane) in cells which retain a long axis and an aspect ratio of >1.2 during mitosis.

Using this system, we were able to describe the process of spindle orientation in detail. Spindles aligned to the plane of the epithelium shortly after NEB, but with a random orientation relative to the long cell axis. Then, over the course of mitosis, spindles rotated within the plane of the epithelium in a process that depended on NuMA/Mud, Dlg and astral microtubules, as previously reported [23]. Through this analysis we also noted that rotating spindles also tended to move off-centre during mitosis in a manner that depended on the same cortical cues, implying that the distribution of cortical forces that act on the two spindle poles to position the spindle are often unbalanced. Interestingly, while important, this aspect of spindle positioning has rarely been included in models of symmetric cell division, which have usually assumed that spindles rotate about the cell centre in response to cortical forces [23,28,37,53]. Therefore to correct spindle off-centring, additional mechanisms will be required to ensure equal daughter cell size at or after, mitotic exit [54–56].

We also found that spindle orientation is a noisy process, since the net angular displacement of the spindle from NEB until anaphase was about half of the cumulative rotation. Despite this, spindles were able to re-orient to the long cell axis on average, such that even though spindle orientation was random at NEB, majority of spindles were closely aligned to the long cell axis by anaphase. Additionally, the extent of spindle rotation in the notum was found to depend on its initial position with respect to the long cell axis. Thus, spindles oriented furthest from the long cell axis at NEB tended to undergo the largest net rotation towards the long cell axis. This implies that spindles experience a torque that depends on the distance from the long cell axis. This stereotypic rotation would be consistent with a model where forces on the spindle were concentrated at opposing ends of the cell, through polarisation of proteins such as Mud/NuMA or Dynein [23,57,58].

How though do spindles read mitotic cell shape? While cell shape, TCJs, and/or anisotropic tissue tension have all been implicated in spindle orientation to the long cell axis, it has been hard to separate their relative contributions to the process. Some studies have pointed to a direct role for actomyosin-based tension on the mitotic cell [29,36,38,57], whereas others have suggested that spindles align with mitotic cell shape polarised by the tension axis [20–22,39]. By comparing the dynamics of spindle orientation in elongated cells inside and outside the midline, where there are marked differences in tissue tension [46], we have been able to demonstrate a role for isotropic tissue tension in long-axis divisions, independent of cell shape. Thus, whereas spindles rotate dynamically to orient with the long axis of cells outside of the midline, they are less effective at doing so within the crowded tissue midline, where cells are very elongated despite being under little if any tissue tension. In line with this, the correlation between spindle orientation and cell elongation was poor for cells in the midline. Thus, spindles in the midline do not follow the long-axis rule. Our findings build on previous studies in the notum, which did not separate the effect of region-specific tissue mechanics or the initial orientation of the spindle [23]. Moreover, since long-axis divisions in the midline are poor, these data might help explain why alternative mechanisms for refining cell packing such as neighbour exchange [47] and cell delamination [46] play such an important roles in this tissue.

In support of a role for isotropic tension in long-axis divisions in the notum, we were able to switch these region-specific spindle behaviours by manipulating tissue-wide levels of active non-muscle Myosin II. Thus, conditions that increased isotropic tissue tension rescued the ability of spindles in cells within the midline to rotate persistently to the long cell axis, and led to a strong correlation between cellular aspect ratio and spindle alignment. By contrast, conditions that reduced overall levels of tension in the tissue outside of the midline (i.e. dominant negative constructs, RNAi or laser ablation) compromised the ability of spindles to rotate persistently towards the long cell axis, even though TCJ polarity and cell shape remained unchanged. Furthermore, directional spindle rotation was lost when cells were mechanically isolated from their neighbours in mid-mitosis in tissues with elevated tension. This suggests that tissue tension is able to influence spindle positioning during mitosis, as has been previously described for cells in culture [28,38,51,52]. Moreover, since perturbations in Myosin II activity have a similar impact on spindle dynamics in MDCK monolayers (Lisica et al., unpublished data), this is likely to be a general phenomenon.

In the notum under conditions of low tissue tension, we also noted that spindle rotation remained dynamic. Moreover, spindles were still pulled off-centre under these conditions. This phenotype, in which spindle movements appear jittery and random, differs from that seen when cortical regulators of spindle orientation were silenced using RNAi. These data suggest that cortical force generators remain active in cells in regions of the tissue with little tension, but are not properly integrated across the entire cell. How then does tissue tension promote the persistent rotation of spindles towards the long axis?

Some insights can probably be gained by looking at spindle aligment in single cells in isolation, where the mitotic cortex is mechanosensitive [59], polarised as the result of the local resistence of retraction fibers to mitotic rounding [37,38,57,58,60], and able to act as a physical platform upon which astral microtubules can pull and exert torque on the spindle. Similar rules are likely to apply in the context of an epithelium, where tricellular junctions will resist mitotic rounding in a tissue subject to isotropic tension, generating a mechanically rigid cortex upon which Dynein can pull to orient the spindle. Conversely, in a crowded tissue, the ability of Dynein anchored in a relatively soft cortex to effectively pull on spindle poles to orient the spindle will be compromised. In this way, even though the actomyosin cytoskeleton may not directly influence the activity of local cortical force generators [61], it will have a profound impact on the ability of the spindle to read and integrate these cues in the mitotic cortex. This makes both tissue tension and cortical mechanics key factors in enabling cells to divide along their long cell axes.

## ACKNOWLEDGEMENTS

We would like to thank past and present members of the Baum lab, Charras lab and Mao lab for advice on the project and for their help with the manuscript. We would also like to thank the funders who enabled this work: ML was funded by the Agency for Science, Technology and Research (Singapore) and supported by the Wellcome Trust Developmental and Stem Cell Biology PhD programme. ML, NR and BB received support from the MRC (MC_CF12266). BB and NR thank Cancer Research UK for programme grant support (C1529/A1734). BB also thanks the BBSRC for support (BB/J008532/1, BB/K009001/1) and NR thanks the BBSRC for support (BB/). G.C. is supported by a consolidator grant from the European Research Council (MolCellTissMech, agreement 647186). YM is funded by a Medical Research Council Fellowship MR/L009056/1 and a UCL Excellence Fellowship.

## AUTHOR CONTRIBUTIONS

BB and ML designed the experiments; ML and NR conducted the experiments; ML and AL analysed the data; BB and ML wrote the paper; GC and YM adviced on the experiments and paper.

---

**Fig S1.**
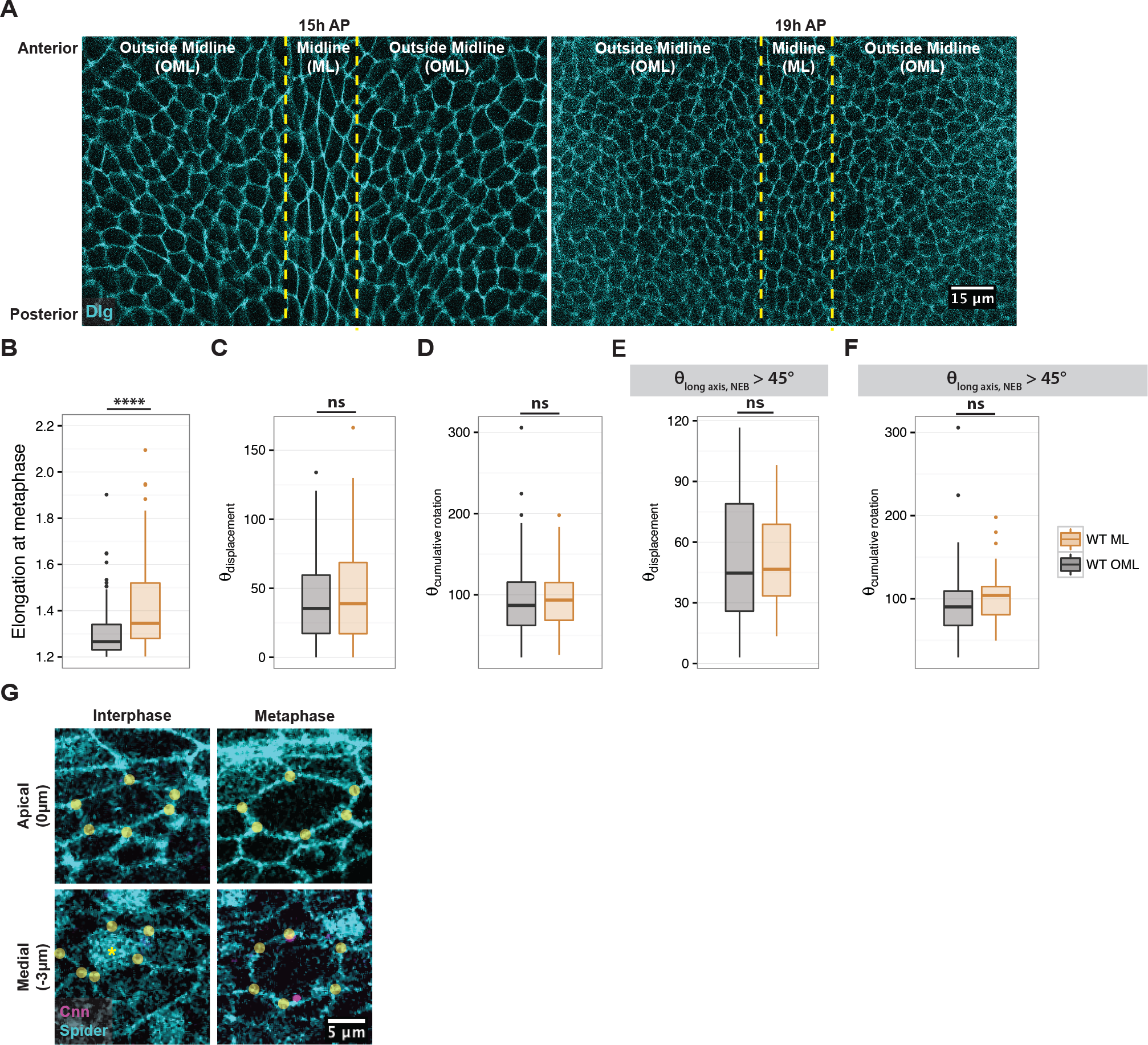
**A**: Image of *Drosophila* pupal notum at 15h and 19h AP. Cell outlines are marked with Dlg-YFP (cyan). Region classified as the midline (ML) are within yellow dotted lines. **B**: Elongation of OML and ML cells at metaphase. Elongation was calculated as the ratio of cell length: width for cells at the height of the spindle (medial plane) at metaphase (1min before anaphase). **C**: Spindle angular displacement (θ_displacement_) for OML and ML spindles. θ_displacement_ was calculated as the angle between the spindle at anaphase and at NEB. θ_displacement_ was similar for both OML and ML spindles (OML: 42.76°±3.48°, n=91; ML: 44.74°±3.89°, n=72). **D**: Cumulative spindle rotation (θ_cumulative rotation_) for OML and ML spindles. θ_cumulative rotation_ was calculated as the sum of the angular distance per min over mitosis. θ_cumulative rotation_ was similar for both OML and ML spindles (OML: 94.55°±4.87 °, n=91; ML: 97.37°±4.66°, n=72). **H**: θ_displacement_ for OML and ML spindles where θ_long axis_ >45° at NEB. θ_displacement_ was similar for OML and ML spindles (OML: 49.42°±4.42°, n=47; ML: 51.00°±4.12°, n=31). **I**: θ_cumulative rotation_ for OML and ML spindles where θ_long axis_ >45° at NEB. θ_cumulative rotation_ was similar for OML and ML spindles (OML: 95.57°±7.04°, n=47; ML: 104.11°±6.29°, n=31) **G**: Distribution of tricellular junctions at apical and medial planes, in an example ML cell before and during mitosis. Cell membranes including nuclear membrane labelled with Spider-GFP (cyan) and spindle poles are labelled with Centrosomin-RFP (magenta). TCJs indicated by yellow dots, pre-NEB nucleus indicated with yellow asterisk. TCJ distribution at apical and medial planes is similar. TCJs are clustered at cell poles, along cell long axis, but the spindle is oriented along cell short axis.

**Fig S2.**
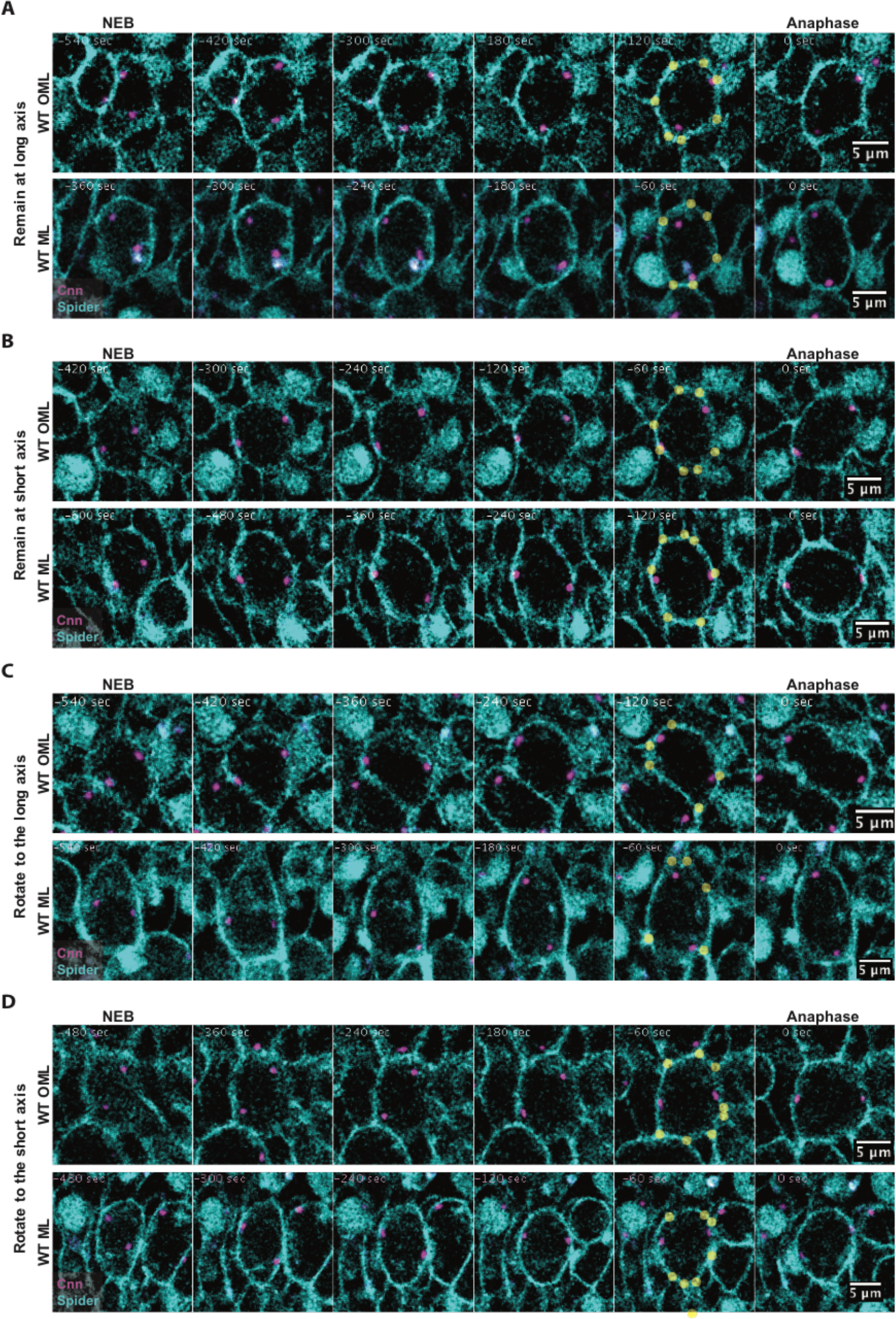
**A**: Spindle orientation from NEB to anaphase over 6 selected timepoints for example OML and ML cell, showing spindles that remain <45° at NEB and anaphase. Cell membranes including nuclear envelope are labelled with Spider-GFP (cyan), spindle poles are labelled with Centrosomin-RFP (magenta) and TCJs are indicated with yellow dots. **B**: Spindle orientation from NEB to anaphase over 6 selected timepoints for example OML and ML cell, showing spindles that remain >45° at NEB and anaphase. Cell membranes including nuclear envelope are labelled with Spider-GFP (cyan), spindle poles are labelled with Centrosomin-RFP (magenta) and TCJs are indicated with yellow dots. **C**: Spindle orientation from NEB to anaphase over 6 selected timepoints for example OML and ML cell, showing spindles that are >45° at NEB but <45° anaphase. Cell membranes including nuclear envelope are labelled with Spider-GFP (cyan), spindle poles are labelled with Centrosomin-RFP (magenta) and TCJs are indicated with yellow dots. **D**: Spindle orientation from NEB to anaphase over 6 selected timepoints for example OML and ML cell, showing spindles that are <45° at NEB but >45° anaphase. Cell membranes including nuclear envelope are labelled with Spider-GFP (cyan), spindle poles are labelled with Centrosomin-RFP (magenta) and TCJs are indicated with yellow dots.

**Fig S3.**
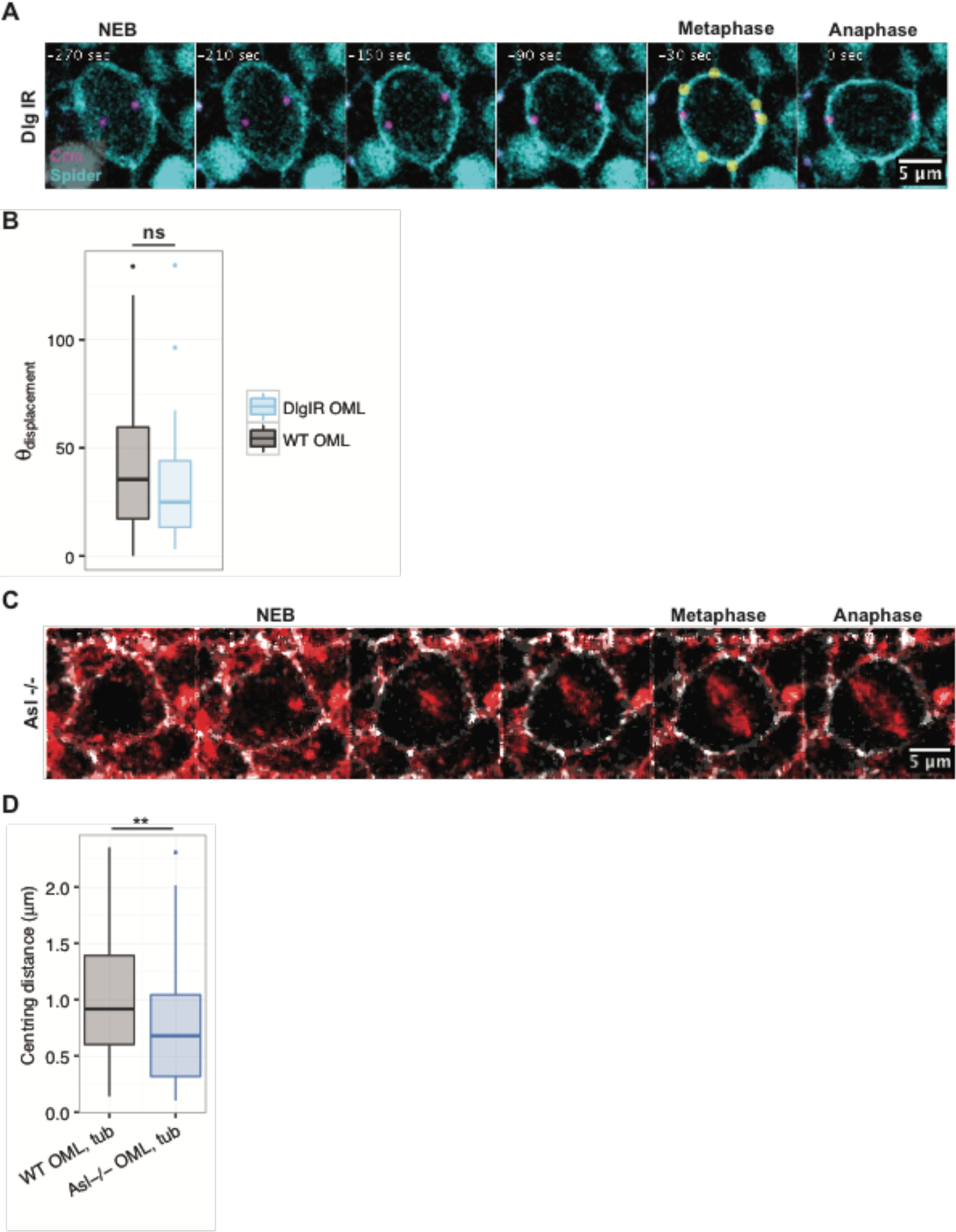
**A**: Example DlgIR OML cell with no *z*-positioning defect and some *xy*-positioning defects during mitosis. Yellow dots indicate TCJs. **B**: θ_displacement_ for WT and DlgIR OML spindles. θ_displacement_ is similar for WT and DlgIR spindles (DlgIR: 32.99°± 4.50°, n=37; p= 0.17). **C**: Example OML cell with Asl^−/−^ mutation and expressing Tubulin-mCherry. *z*-positioning defects are very prevalent in Asl^−/−^ mutants, but a mitotic spindle is still able to form that is occasionally within the plane of the tissue at metaphase (shown). **D**: Centring distance for WT and Asl^−/−^ OML spindles labelled with Tubulin-mCherry. Asl^−/−^spindles were significantly closer to the centre of the cell compared to WT spindles (WT: 1.00±0.07, n= 58; Asl: 0.76±0.05, n=68, p=0.0090).

**Fig S4.**
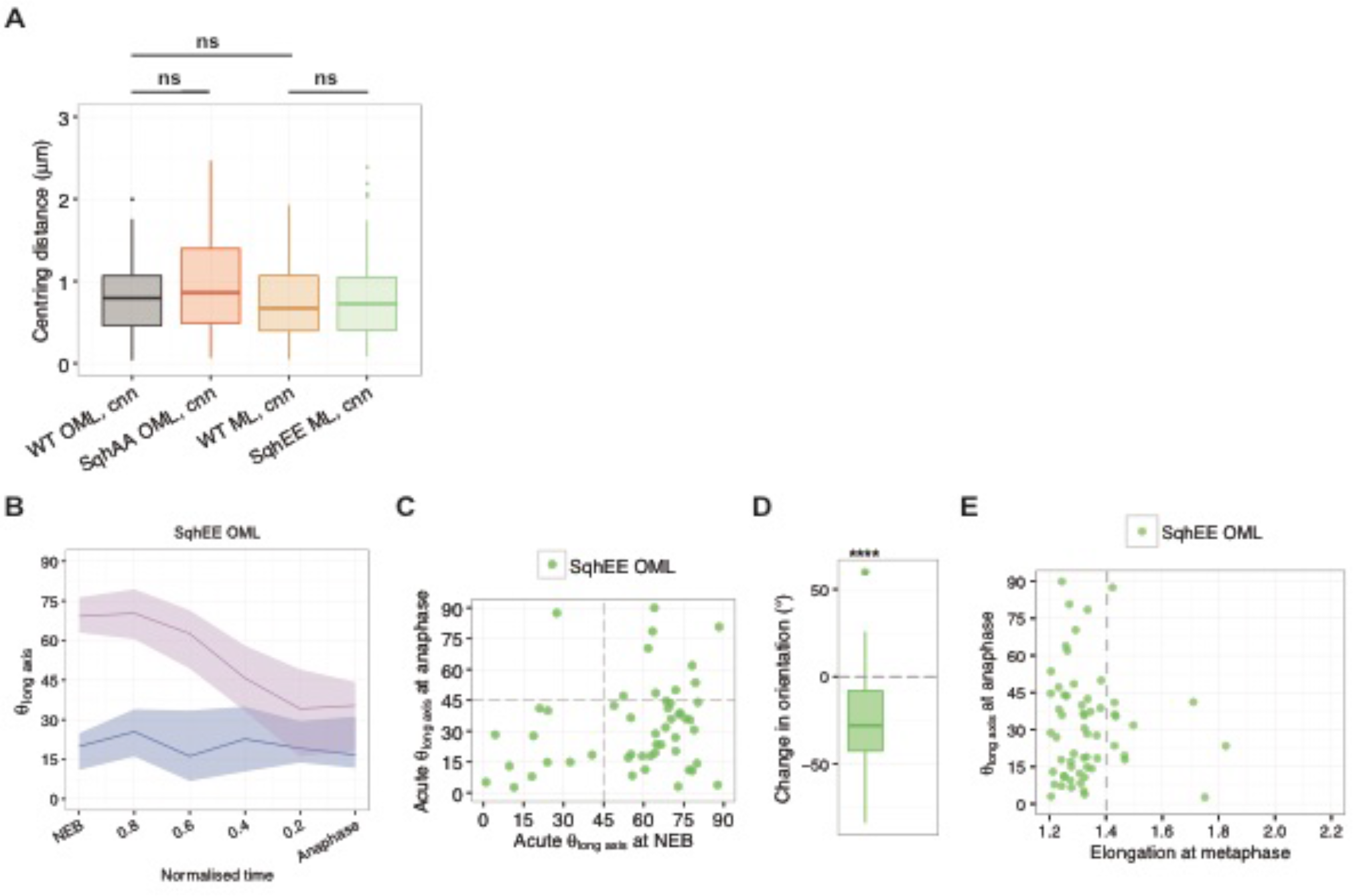
**A**: Centring distance for WT OML, SqhAA OML, WT ML and SqhEE ML spindles labelled with Cnn-RFP. Centring distance is similar between experimental conditions. **B**: θ_long axis_ from NEB to anaphase for SqhEE OML spindles. Data is shown for subpopulation of spindles that are ≥45° (pink line) or <45° (purple line) at NEB. Lines indicate median values and shaded regions indicate interquartile range. **C**: Orientation of spindles at anaphase against orientation at NEB for SqhEE OML spindles. Majority of SqhEE OML spindles were oriented >45° at NEB but <45° at anaphase, with almost none oriented <45° at NEB but >45° at anaphase. **D**: Change in orientation for SqhEE OML spindles. Change in orientation was significantly <0 for SqhEE OML spindles, and less than that of WT OML spindles (SqhEE ML: −24.61± 3.95, p= 6.75×10^−7^, n=51). **E**: Spindle orientation at anaphase against cell elongation at metaphase for SqhEE OML cells. All SqhEE ML cells with higher elongations were oriented <45° at anaphase.

**Fig S5.**
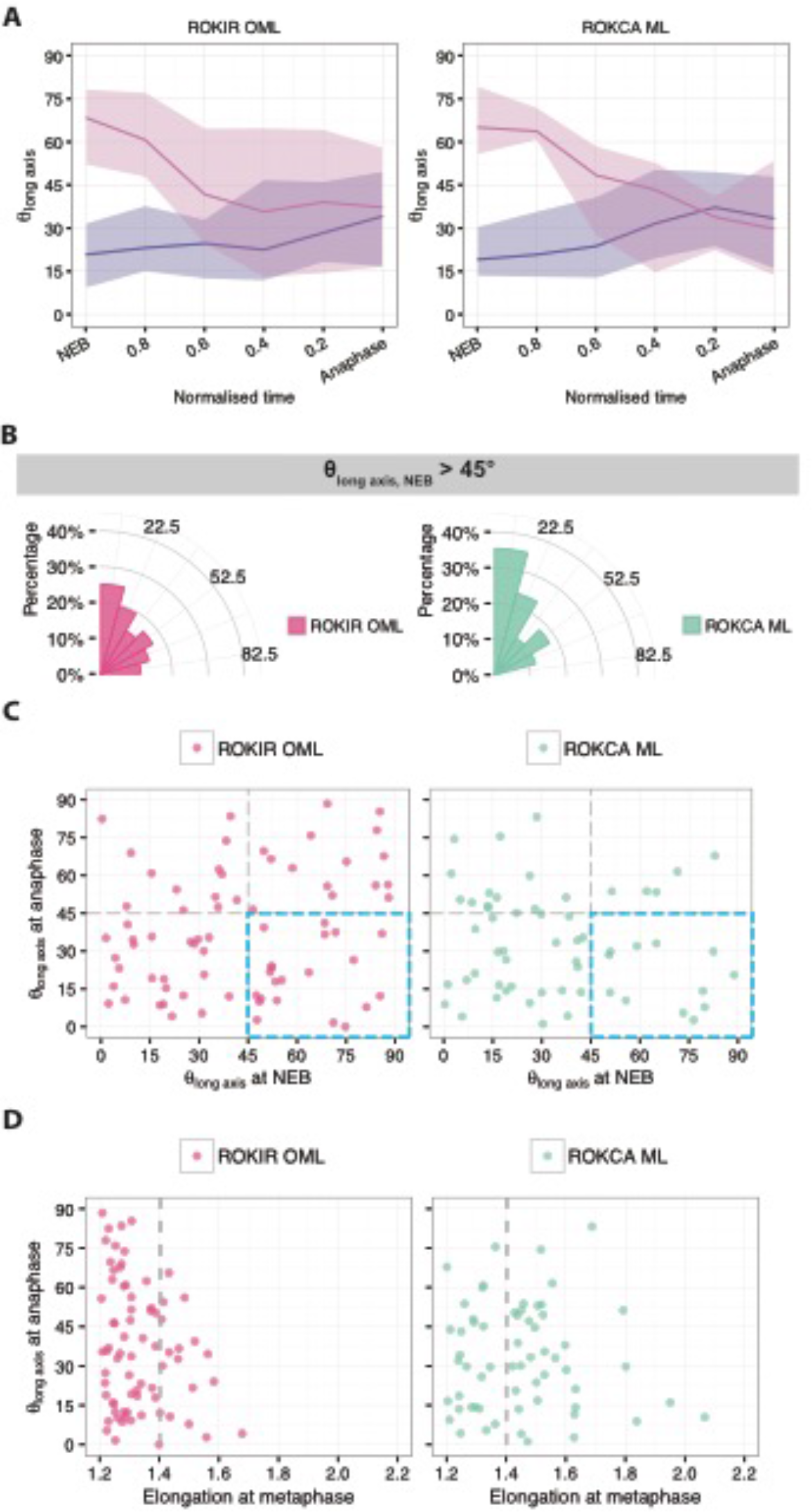
**A**: θ_long axis_ from NEB to anaphase for ROK-IR OML and ROK-CA ML spindles. Data is shown for subpopulation of spindles that are ≥45° (pink line) or <45° (purple line) at NEB. Lines indicate median values and shaded regions indicate interquartile range. **B**: θ_long axis_ at anaphase for ROK-IR OML and ROK-CA ML spindles that were >45° at NEB. The distribution of θ_long axis_ for ROK-IR OML spindles is wider than for WT OML spindles, the distribution of θ_long axis_ for ROK-CA ML spindles is narrower and closer to 0° than WT ML spindles. **C**: Orientation of spindles at anaphase against at NEB for ROK-IR OML and ROK-CA ML spindles. Blue box highlights spindles that are oriented >45° at NEB but <45° at anaphase. **D**: Spindle orientation at anaphase against cell elongation at metaphase for ROK-IR OML and ROK-CA ML cells. ROK-IR OML cells at higher elongations (>1.4) had orientations of >45° at anaphase, while almost all ROK-CA ML cells with higher elongations were oriented <45° at anaphase.

## METHODS

### Live-imaging

*Drosophila* pupae were selected at the white pre-pupal stage, 0h after pupariation (AP), and imaged at 14.5h AP for 2-3h at room temperature. Developmental time was halved when incubated at 29°C; and doubled when incubated at 18°C. Pupae for live imaging were attached to a glass slide ventral side down with double-sided tape between spacers made with small glass coverslips. The pupal case was removed from the dorsal side of the animal and a glass coverslip coated with mineral oil on one side was placed over the spacers, just touching the dorsal tissue of the pupa. The entire set-up was placed under the microscope for live-imaging (Zitserman and Roegiers, 2011; Georgiou and Baum, 2010).

Imaging was done on Leica SPE and SP5 confocal microscopes with a 63X lens (N.A.) or 60X lens (N.A.) respectively.

### Immunohistochemistry

*Drosophila* pupae for immunostaining were dissected at 15h AP. Pupae were pinned with sharpened wires dorsal side down onto a PDMS dish filled with PBS. The pupal head was removed with small surgical scissors and the ventral length of the pupa was cut out. The dorsal tissue around the notum was isolated and transferred into glass wells with micropippettes for fixing and staining. Dissected nota were stained with the following antibodies and probes:

| Antibody/ probe | Concentration | Source |
| --- | --- | --- |
| Mouse anti-Discs large | 1: 100 | DSHB |

Alexa conjugated fluorophores were used in secondary stains. Immunostained nota were imaged on Leica SPE confocal microscopes with a 63X lens (N.A.)

### Fly stocks used

#### BACKGROUND AND VISUALISING OF CELL OUTLINE AND MITOTIC STRUCTURES

| Stock | Source |
| --- | --- |
| w<sup>1118</sup>;;; | BL 3605 |
| w<sup>1118</sup>;; pnr-GAL4; | BL 3039 |
| ;; Spider-GFP;; | Flytrap Insertion<br>(BL 59025) |
| w<sup>1118</sup>;; Spider-GFP, pnr-GAL4; | Recombined strain of BL 3039 and<br>BL 59025 by Rodriguez, N. |
| Discs large (Dlg)-YFP;;; | Cambridge Protein Trap Insertion<br>(DGRC 115375) |
| w<sup>1118</sup>; ubi-RFP-Cnn;;; | Raff, J. |
| ; actin-GAL4, UAS-mCherry-α-Tubulin;;; | Recombined from BL4414 and<br>BL25774 by Rodrigues N. |
| w*; UAS-mCherry-α-Tubulin;;; | BL 25774 |

#### RNAI-MEDIATED SILENCING

Interfering RNA transcripts targeting expression of proteins were expressed using the GAL4/ UAS system [1]. GAL4 expression was under the control of the *pannier* gene (Pnr-GAL4) [2], restricting GAL4 binding of UAS response elements and subsequent expression of constructs to the central region of the notum. Pupae in RNAi experiments were incubated at 25°C or 29°C from 9-14.5h AP or 0-14.5h AP to ensure efficient expression of GAL4. Where lethality was seen under these conditions, pupae were incubated at 18°C from 0-14.5h AP to reduce the activity of GAL4.

The following fly stocks were used in RNAi-mediated silencing:

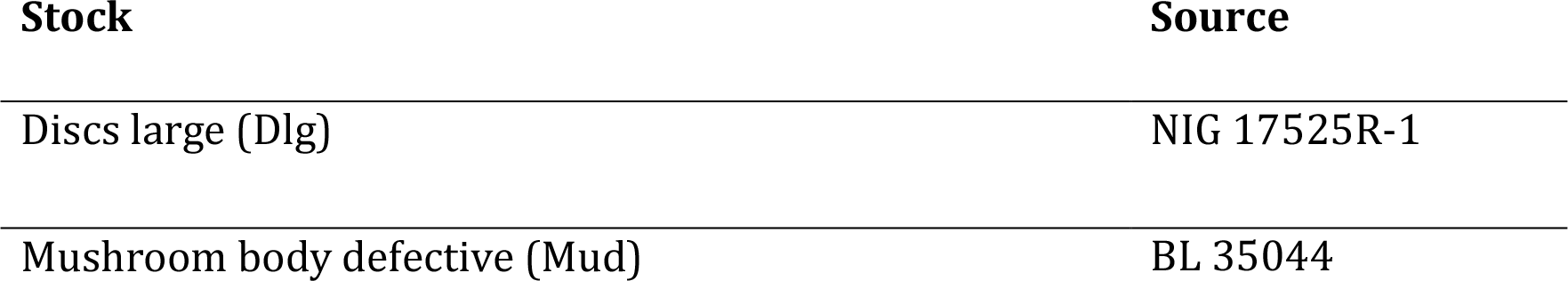

#### PROTEIN CODING CONSTRUCTS

Constructs were expressed using the GAL4/ UAS system [1]. GAL4 expression was under the control of the *pannier* gene (Pnr-GAL4) [2], restricting GAL4 binding of UAS response elements and subsequent expression of constructs to the central region of the notum. All pupae were incubated at 25°C or 29°C from 9-14.5h AP or 0-14.5h AP to ensure efficient expression of GAL4.

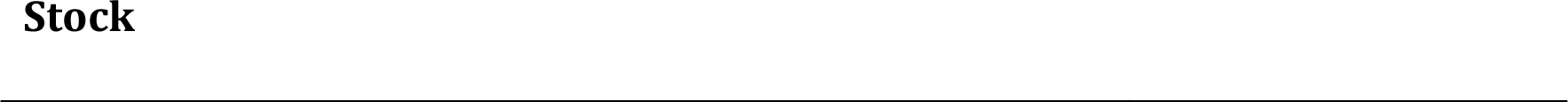
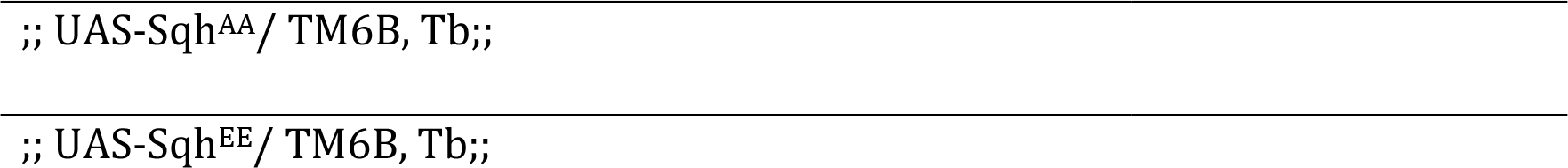

#### ASTERLESS^MECD^ EXPERIMENTS

;; Asl^mecD^/ TM6B, Tb;; flies were a gift from Raff, J. Experiments were done by crossing Dlg-YFP;; AslmecD/ TM6B, Tb;; flies with; actin-GAL4-UAS-mCherry-α-Tubulin; Asl^mecD^/ TM6B, Tb;; flies. Pupae homozygous for the Asl^mecD^ allele were identified by selecting against TM6, Tb (tubby pupae).

### Mechanical isolation by laser ablaton

w^1118^; ubi-RFP-Cnn; Spider-GFP, Pnr-Gal4/ UAS-Sqh^EE^;; pupae were prepared for live-imaging and mounted on an inverted Zeiss multiphoton. Cells about to undergo mitosis were visually identified by the presence of centrosomes around the nucleus. The region to be ablated was manually marked at the apical surface, and irradiated with a UV laser. Z-stack images were acquired in Airyscan mode, every 15 seconds. Ablation was carried out after the first frame, and imaging continued until anaphase was observed.

### Image analysis

#### QUANTIFICATION OF CELL SHAPE

The medial plane was identified as the plane where majority of the spindle was located, which was usually the plane with both spindle poles visible. The cell outline in the medial plane was manually marked out in FIJI (http://fiji.sc/Fiji). The centroid of the outline was taken as the cell centre, while the major length and minor length of the fit ellipse to the outline were taken as length and width of the cell. The angle of the major length of the fit ellipse was taken as the orientation of the long axis of the cell.

#### QUANTIFICATION OF SPINDLE MOVEMENTS

Spindle movement was tracked by drawing a line between the visible spindle poles from NEB through to anaphase. Spindle angle, centroid and length were recorded, and spindle pole coordinates were calculated from these values. Spindle pole coordinates for a
spindle with centroid coordinates (*x*,*y*), length *l* and angle *θ*, were calculated as (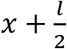.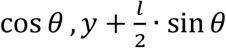) and (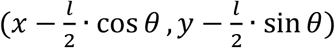). Spindles were not considered for analysis if apparent spindle poles were more than 1.5 μm apart. Spindle measurements were taken with Tubulin-mCherry marking the spindle or Centrosomin-RFP marking the spindle poles. Calculations of spindle angles were similar for measurements done with Tubulin or Centrosomin as a marker, and so the results were pooled. Calculations of spindle length and centroid (including spindle pole coordinates) were significantly different when comparing between measurements taken with Tubulin or Centrosomin as a marker. This was likely due to tubulin being less precise for identifying the spindle poles and therefore spindle length and centre. These measurements were separated based on markers across perturbations, and the markers used in each analysis are identified in the text.

#### IDENTIFICATION OF MITOTIC EVENTS

Mitotic time was taken as the time from nuclear envelope breakdown (NEB) till anaphase onset. NEB was identified as the first timeframe when nuclear exclusion of background fluorescent signal disappeared. Anaphase onset was identified in cells expressing Tubulin-mCherry as the first timeframe when tubulin accumulation towards the spindle poles was observed, ~ 3min before furrow ingression begins. In movies using only Centrosomin-RFP as a marker, anaphase onset was taken as the timeframe 3min before furrow ingression begins. Late metaphase was defined as 1min before anaphase onset.

### Statistical analysis and data visualisation

Two sample Wilcoxon ranked sum test was performed to compare medians between data, using the wilcox.test() function in R. Two sample Kolmogorov-Smirnov test was performed to compare distributions of datasets, using the ks.test() function in R. Random uniform distributions were generated with the runif() function in R.

Graphpad Prism and R (ggplot2 library) were used to generate graphs representing the data. Line plots representing median over time, with error bars representing interquartile range were generated in Prism. Individual line plots representing spindle orientation over time were generated in R. Boxplots were generated with the geom_boxplot() function in R, with the boxes representing the upper quartile, median and lower quartile of the data, and whiskers representing the data within 1.5 times the interquartile range flanking the upper and lower quartiles. All remaining data (outliers) are represented as points. All data was plotted, unless stated otherwise in the text. Linear regressions were performed with the lm() and geom_smooth(method=lm) function in R, and best-fit linear mean lines with standard error of the mean were plotted. All stated R2 values are adjusted R2 values.

## REFERENCES

1. Bilder, D., Li, M., Perrimon, N., Bilder, D., Perrimon, N., Harrison, D.A., Perrimon, N., Jacob, L., Opper, M., Metzroth, B., et al. (2000). Cooperative regulation of cell polarity and growth by Drosophila tumor suppressors. Science 289, 113–6.

2. Laprise, P., and Tepass, U. (2011). Novel insights into epithelial polarity proteins in Drosophila. Trends Cell Biol. 21, 401–8.

3. Rodriguez-Boulan, E., and Macara, I.G. (2014). Organization and execution of the epithelial polarity programme. Nat. Rev. Mol. Cell Biol. 15, 225–242.

4. Morin, X., and Bellaïche, Y. (2011). Mitotic spindle orientation in asymmetric and symmetric cell divisions during animal development. Dev. Cell 21, 102–19.

5. di Pietro, F., Echard, A., and Morin, X. (2016). Regulation of mitotic spindle orientation: an integrated view. EMBO Rep. 17, 1106–1130.

6. Rappaport, R. (1997). Cleavage furrow establishment by the moving mitotic apparatus. Dev. Growth Differ. 39, 221–226.

7. White, E.A., and Glotzer, M. (2012). Centralspindlin: At the heart of cytokinesis. Cytoskeleton 69, 882–892.

8. Williams, S.E., and Fuchs, E. (2013). Oriented divisions, fate decisions. Curr. Opin. Cell Biol.

9. Luxenburg, C., Pasolli, H.A., Williams, S.E., and Fuchs, E. (2011). Developmental roles for Srf, cortical cytoskeleton and cell shape in epidermal spindle orientation. Nat. Cell Biol. 13, 203–14.

10. Nakajima, Y.-I., Kuranaga, E., Sugimura, K., Miyawaki, A., and Miura, M. (2011). Nonautonomous apoptosis is triggered by local cell cycle progression during epithelial replacement in Drosophila. Mol. Cell. Biol. 31, 2499–512.

11. Padash-Barmchi, M., Charish, K., Que, J., and Auld, V.J. (2013). Gliotactin and Discs Large Are Co-Regulated To Maintain Epithelial Integrity. J. Cell Sci. 126, 1134–43.

12. Poulton, J.S., Cuningham, J.C., and Peifer, M. (2014). Acentrosomal Drosophila Epithelial Cells Exhibit Abnormal Cell Division, Leading to Cell Death and Compensatory Proliferation. Dev. Cell 30, 731–745.

13. Ganier, O., Schnerch, D., Oertle, P., Lim, R.Y., Plodinec, M., and Nigg, E.A. (2018). Structural centrosome aberrations promote non-cell-autonomous invasiveness. EMBO J., e98576.

14. Cadart, C., Zlotek-Zlotkiewicz, E., Le Berre, M., Piel, M., and Matthews, H.K. (2014). Exploring the Function of Cell Shape and Size during Mitosis. Dev. Cell 29, 159–169.

15. Kiyomitsu, T. (2015). Mechanisms of daughter cell-size control during cell division. Trends Cell Biol. 25, 286–95.

16. Hertwig, O. (1896). The Biological Problem of To-day Preformation or epigenesis? The basis of a theory of organic development Translated.. W. Heinemann, ed. (London: Heinemann’s Scientific Handbooks)

17. Gibson, W.T., and Gibson, M.C. (2009). Cell topology, geometry, and morphogenesis in proliferating epithelia. Curr. Top. Dev. Biol. 89, 87–114.

18. Minc, N., and Piel, M. (2012). Predicting division plane position and orientation. Trends Cell Biol. 22, 193–200.

19. Mao, Y., Tournier, A.L., Hoppe, A., Kester, L., Thompson, B.J., and Tapon, N. (2013). Differential proliferation rates generate patterns of mechanical tension that orient tissue growth. EMBO J. 32, 2790–2803.

20. Campinho, P., Behrndt, M., Ranft, J., Risler, T., Minc, N., and Heisenberg, C.-P. (2013). Tension-oriented cell divisions limit anisotropic tissue tension in epithelial spreading during zebrafish epiboly. Nat. Cell Biol. advance on.

21. Mao, Y., Tournier, A.L., Bates, P.A., Gale, J.E., Tapon, N., and Thompson, B.J. (2011). Planar polarization of the atypical myosin Dachs orients cell divisions in Drosophila. Genes Dev. 25, 131–6.

22. Wyatt, T.P.J., Harris, A.R., Lam, M., Cheng, Q., Bellis, J., Dimitracopoulos, A., Kabla, A.J., Charras, G.T., and Baum, B. (2015). Emergence of homeostatic epithelial packing and stress dissipation through divisions oriented along the long cell axis. Proc. Natl. Acad. Sci. U. S. A. 112, 5726–31.

23. Bosveld, F., Markova, O., Guirao, B., Martin, C., Wang, Z., Pierre, A., Balakireva, M., Gaugue, I., Ainslie, A., Christophorou, N., et al. (2016). Epithelial tricellular junctions act as interphase cell shape sensors to orient mitosis. Nature.

24. Grill, S.W., and Hyman, A.A. (2005). Spindle positioning by cortical pulling forces. Dev. Cell 8, 461–5.

25. Kotak, S., and Gönczy, P. (2013). Mechanisms of spindle positioning: cortical force generators in the limelight. Curr. Opin. Cell Biol. 16, 1–8.

26. Wühr, M., Tan, E.S., Parker, S.K., Detrich, H.W., Mitchison, T.J., Dogterom, M., Kerssemakers, J.W., Romet-Lemonne, G., Janson, M.E., Grill, S.W., et al. (2010). A model for cleavage plane determination in early amphibian and fish embryos. Curr. Biol. 20, 2040–5.

27. Kimura, K., and Kimura, A. (2011). A novel mechanism of microtubule length-dependent force to pull centrosomes toward the cell center. Bioarchitecture 1, 74–79.

28. Minc, N., Burgess, D., and Chang, F. (2011). Influence of cell geometry on division-plane positioning. Cell 144, 414–26.

29. Gloerich, M., Bianchini, J.M., Siemers, K.A., Cohen, D.J., and Nelson, W.J. (2017). Cell division orientation is coupled to cell–cell adhesion by the E-cadherin/LGN complex. Nat. Commun. 8, 13996.

30. Dujardin, D.L., and Vallee, R.B. (2002). Dynein at the cortex. Curr. Opin. Cell Biol. 14, 44–49.

31. Kraft, L.M., and Lackner, L.L. (2017). Mitochondria-driven assembly of a cortical anchor for mitochondria and dynein. J. Cell Biol. 216, 3061–3071.

32. Radulescu, A.E., and Cleveland, D.W. (2010). NuMA after 30 years: the matrix revisited. Trends Cell Biol. 20, 214–222.

33. Heppert, J.K., Pani, A.M., Roberts, A.M., Dickinson, D.J., and Goldstein, B. (2018). A CRISPR tagging-based screen reveals localized players in wnt-directed asymmetric cell division. Genetics 208, 1147–1164.

34. Dimitracopoulos, A., Lam, M., and Baum, B. (2016). Oriented Division: Using T-Junctions to Determine Direction. Curr. Biol. 26, R371–R373.

35. Théry, M., Racine, V., Pépin, A., Piel, M., Chen, Y., Sibarita, J.-B., and Bornens, M. (2005). The extracellular matrix guides the orientation of the cell division axis. Nat. Cell Biol. 7, 947–53.

36. Théry, M., Racine, V., Piel, M., Pepin, A., Dimitrov, A., Chen, Y., Sibarita, J.-B., and Bornens, M. (2006). Anisotropy of cell adhesive microenvironment governs cell internal organization and orientation of polarity. Proc. Natl. Acad. Sci. 103, 19771–19776.

37. Théry, M., Jiménez-Dalmaroni, A., Racine, V., Bornens, M., and Jülicher, F. (2007). Experimental and theoretical study of mitotic spindle orientation. Nature 447, 493–6.

38. Fink, J., Carpi, N., Betz, T., Bétard, A., Chebah, M., Azioune, A., Bornens, M., Sykes, C., Fetler, L., Cuvelier, D., et al. (2011). External forces control mitotic spindle positioning. Nat. Cell Biol. 13, 771–8.

39. Nestor-bergmann, A., Stooke-vaughan, G.A., Goddard, G.K., Starborg, T., Jensen, O.E., and Woolner, S. (2017). Decoupling the roles of cell shape and mechanical stress in orienting and cueing epithelial mitosis.

40. Tang, Z., Hu, Y., Wang, Z., Jiang, K., Zhan, C., Marshall, W.F., and Tang, N. (2018). Mechanical Forces Program the Orientation of Cell Division during Airway Tube Morphogenesis. Dev. Cell 44, 313–325.e5.

41. Bonnet, I., Marcq, P., Bosveld, F., Fetler, L., Bellaïche, Y., and Graner, F. (2012). Mechanical state, material properties and continuous description of an epithelial tissue. J. R. Soc. Interface 9, 2614–23.

42. Bosveld, F., Bonnet, I., Guirao, B., Tlili, S., Wang, Z., Petitalot, A., Marchand, R., Bardet, P.-L., Marcq, P., Graner, F., et al. (2012). Mechanical Control of Pathway. Science.

43. Rosa, A., Vlassaks, E., Pichaud, F., and Baum, B. (2015). Ect2/Pbl Acts via Rho and Polarity Proteins to Direct the Assembly of an Isotropic Actomyosin Cortex upon Mitotic Entry. Dev. Cell, 604–616.

44. Rodrigues, N.T.L., Lekomtsev, S., Jananji, S., Kriston-Vizi, J., Hickson, G.R.X., and Baum, B. (2015). Kinetochore-localized PP1–Sds22 couples chromosome segregation to polar relaxation. Nature 524, 489–492.

45. Hunter, G.L., Hadjivasiliou, Z., Bonin, H., He, L., Perrimon, N., Charras, G., and Baum, B. (2016). Coordinated control of Notch/Delta signalling and cell cycle progression drives lateral inhibition-mediated tissue patterning. Development 143, 2305–10.

46. Marinari, E., Mehonic, A., Curran, S., Gale, J., Duke, T., and Baum, B. (2012). Live-cell delamination counterbalances epithelial growth to limit tissue overcrowding. Nature 484, 542–5.

47. Curran, S., Strandkvist, C., Bathmann, J., de Gennes, M., Kabla, A., Salbreux, G., and Baum, B. (2017). Myosin II Controls Junction Fluctuations to Guide Epithelial Tissue Ordering. Dev. Cell 43, 480–492.e6.

48. Pecreaux, J., Röper, J.-C., Kruse, K., Jülicher, F., Hyman, A.A., Grill, S.W., Howard, J., Grill, S.W., Kruse, K., Julicher, F., et al. (2006). Spindle Oscillations during Asymmetric Cell Division Require a Threshold Number of Active Cortical Force Generators. Curr. Biol. 16, 2111–2122.

49. Izumi, Y., Ohta, N., Hisata, K., Raabe, T., and Matsuzaki, F. (2006). Drosophila Pins-binding protein Mud regulates spindle-polarity coupling and centrosome organization. Nat. Cell Biol. 8, 586–93.

50. Kiyomitsu, T., and Cheeseman, I.M. (2012). Chromosome- and spindle-pole-derived signals generate an intrinsic code for spindle position and orientation. Nat. Cell Biol. 14, 311–317.

51. O’Connell, C.B., and Wang, Y.-l. (2000). Mammalian Spindle Orientation and Position Respond to Changes in Cell Shape in a Dynein-dependent Fashion. Mol. Biol. Cell 11, 1765–1774.

52. Fernandez, P., Maier, M., Lindauer, M., Kuffer, C., Storchova, Z., and Bausch, A.R. (2011). Mitotic spindle orients perpendicular to the forces imposed by dynamic shear. PLoS One 6, e28965.

53. Gibson, W.T., Veldhuis, J.H., Rubinstein, B., Cartwright, H.N., Perrimon, N., Brodland, G.W., Nagpal, R., and Gibson, M.C. (2011). Control of the mitotic cleavage plane by local epithelial topology. Cell 144, 427–38.

54. Tzur, A., Kafri, R., LeBleu, V.S., Lahav, G., and Kirschner, M.W. (2009). Cell growth and size homeostasis in proliferating animal cells. Science (80-.). 325, 167–171.

55. Sung, Y., Tzur, A., Oh, S., Choi, W., Li, V., Dasari, R.R., Yaqoob, Z., and Kirschner, M.W. (2013). Size homeostasis in adherent cells studied by synthetic phase microscopy. Proc. Natl. Acad. Sci. 110, 16687–16692.

56. Kiyomitsu, T., and Cheeseman, I.M. (2013). Cortical Dynein and asymmetric membrane elongation coordinately position the spindle in anaphase. Cell 154, 391–402.

57. Machicoane, M., de Frutos, C.A., Fink, J., Rocancourt, M., Lombardi, Y., Gare, S., Piel, M., and Echard, A. (2014). SLK-dependent activation of ERMs controls LGN-NuMA localization and spindle orientation. J. Cell Biol. 205, 791–9.

58. Matsumura, S., Kojidani, T., Kamioka, Y., Uchida, S., Haraguchi, T., Kimura, A., and Toyoshima, F. (2016). Interphase adhesion geometry is transmitted to an internal regulator for spindle orientation via caveolin-1. Nat. Commun. 7, ncomms11858.

59. Schiffhauer, E.S., Luo, T., Mohan, K., Srivastava, V., Qian, X., Griffis, E.R., Iglesias, P.A., and Robinson, D.N. (2016). Mechanoaccumulative Elements of the Mammalian Actin Cytoskeleton. Curr. Biol.

60. Oakes, P.W., Banerjee, S., Marchetti, M.C., and Gardel, M.L. (2014). Geometry Regulates Traction Stresses in Adherent Cells. Biophys. J. 107, 825–833.

61. Redemann, S., Pecreaux, J., Goehring, N.W., Khairy, K., Stelzer, E.H.K., Hyman, A.A., and Howard, J. (2010). Membrane invaginations reveal cortical sites that pull on mitotic spindles in one-cell C. elegans embryos. PLoS One 5, e12301.

## References

1. Brand, A., and Perrimon, N. (1993). Targeted gene expression as a means of altering cell fates and generating dominant phenotypes. Development 118, 401–415.

2. Calleja, M., Herranz, H., Estella, C., Casal, J., Lawrence, P., Simpson, P., and Morata, G. (2000). Generation of medial and lateral dorsal body domains by the pannier gene of Drosophila. Development 127, 3971–80.

